# Interaction with TopBP1 mediates human papillomavirus 16 E2 plasmid segregation/retention function and stability during the viral life cycle

**DOI:** 10.1101/2022.01.28.478274

**Authors:** Apurva T. Prabhakar, Claire D. James, Dipon Das, Christian T. Fontan, Raymonde Otoa, Xu Wang, Molly L. Bristol, Iain M. Morgan

## Abstract

Human papillomavirus 16 (HPV16) E2 is a DNA binding protein that regulates transcription, replication and potentially, segregation of the HPV16 genome during the viral life cycle. In the segregation model, E2 simultaneously binds to viral and host chromatin, acting as a bridge to ensure that viral genomes reside in daughter nuclei following cell division. The host chromatin receptor for E2 mediating this function is unknown. Recently, we demonstrated that CK2 phosphorylation of E2 on serine 23 (S23) is required for interaction with TopBP1, and that this interaction promotes E2 and TopBP1 recruitment to mitotic chromatin. Here, we demonstrate that in U2OS and N/Tert-1 cells expressing wild type E2 and a non-TopBP1 binding mutant (S23A, serine 23 mutated to alanine), interaction with TopBP1 is essential for E2 recruitment of plasmids to mitotic chromatin. Using a novel quantitative segregation assay, we demonstrate that interaction with TopBP1 is required for E2 plasmid segregation. The interaction of E2 with TopBP1 promotes increased levels of E2 protein during mitosis in U2OS and N/Tert-1 cells, as well as in human foreskin keratinocytes immortalized by the HPV16 genome. Additionally, interaction with TopBP1 is required for expression of the E2 protein during the viral life cycle. Overall, our results demonstrate that E2 has plasmid segregation activity, and that the E2-TopBP1 interaction is essential for this E2 function and for E2 expression during the viral life cycle.

**Importance:** HPV16 causes 3-4% of all human cancers. During the viral life cycle, it is proposed that the viral genome is actively segregated into daughter nuclei, ensuring viral replication in the subsequent S phase. The E2 protein potentially bridges the viral and host genomes during mitosis to mediate segregation of the circular viral plasmid. Here, we demonstrate that E2 has the ability to mediate plasmid segregation, and that this function is dependent upon interaction with the host protein TopBP1. Additionally, we demonstrate that the E2-TopBP1 interaction promotes enhanced E2 expression during mitosis which likely promotes the plasmid segregation function of E2. Overall, our results present a mechanism of how HPV16 can segregate its viral genome during an active infection, a critical aspect of the viral life cycle.

## Introduction

Human papillomaviruses are causative agents in around 5% of all human cancers, targeting anogenital and oropharyngeal regions, with HPV16 being the most prevalent type detected (1). There are no drugs for the treatment of HPV cancers that directly target a viral process, an enhanced understanding of the viral life cycle is required in order to identify novel therapeutic targets. HPVs infect basal epithelial cells and following cell division the viral genomes enter the host nuclei, a cellular compartment required for replication of the double stranded DNA viral genome (2-4). Following entrance into the infected cell nucleus, host factors activate transcription from the viral genome (5). The E6 and E7 viral proteins target and degrade p53 and pRb (among other targets), respectively, promoting cellular proliferation and entry into S phase (6).

During S phase, two viral proteins (E1 and E2) interact with host factors to replicate the viral genome (7, 8). The carboxyl-terminal domain of E2 forms homo-dimers and binds to 12bp palindromic target sequences on the viral long control region (LCR), which controls transcription and replication of the viral genome. Three of the E2 target sequences surround the viral origin of replication and following binding, E2 recruits the E1 protein to the origin of replication via a protein-protein interaction (9, 10). E1 then interacts with a number of cellular replication factors in order to replicate the viral genome (11-16). In addition to promoting viral replication, E2 has additional functions during the viral life cycle. E2 can regulate transcription from the viral genome, activating or repressing depending upon the levels of E2 protein (17). E2 can also regulate transcription from the host genome that has direct relevance to the viral life cycle (18, 19). The final proposed function for E2 during the viral life cycle is to act as a viral segregation factor (20). During mitosis, there is an active mechanism to retain the viral genome in resulting daughter nuclei. It is proposed that E2 acts as a bridge between host and viral genome to ensure localization of the viral genomes in daughter nuclei following cell division. For BPV1 E2, BRD4 is the mitotic chromatin receptor (21-24). However, for high-risk HPV types (those that cause cancer, including HPV16), E2 and BRD4 do not co-localize on mitotic chromatin suggesting that BRD4 is not the mitotic chromatin receptor for these HPV E2 proteins (25, 26).

We identified an interaction between HPV16 E2 (E2 from now on will indicate HPV16) and TopBP1 (27, 28). This interaction is involved in regulating E1-E2 replication function (29-31). TopBP1 is also a strong candidate for mediating the plasmid segregation function for E2. TopBP1 is a multi-functional protein encoding 9 BRCT domains, and is involved in all aspects of nucleic acid metabolism (32). TopBP1 is also highly functional during mitosis as it can prevent transmission of damaged and under-replicated DNA to daughter cells (33-40). Furthermore, TopBP1 regulates the ability of E2 to interact with interphase chromatin, and co-localizes with E2 on mitotic chromatin (41). Recently, we demonstrated that phosphorylation of E2 on serine 23 promotes a direct interaction between these two proteins *in vitro* and *in vivo*, and that E2 recruits TopBP1 onto mitotic chromatin (42). Our work showed that mutation of serine 23 to alanine (S23A) resulted in a compromised interaction of E2 with mitotic chromatin, and that E2-TopBP1 interaction is essential for the HPV16 life cycle (42).

Building on this, we report here that E2 recruits plasmid DNA to mitotic chromatin in a TopBP1 interaction dependent manner in U2OS cells. We describe a novel quantitative assay demonstrating that the E2-TopBP1 interaction is required for E2 segregation/retention function in both U2OS cells, and the human foreskin keratinocyte cell line N/Tert-1. We also demonstrate that in U2OS, N/Tert-1 and human foreskin keratinocytes immortalized by HPV16 (HFK+HPV16), the interaction between E2 and TopBP1 is required for increasing both E2 and TopBP1 protein levels during mitosis. While additional host proteins may contribute to the plasmid segregation function of E2, we have demonstrated that the E2-TopBP1 interaction is critical for this E2 function. In the HPV16 life cycle E2 expression is dependent upon interaction with TopBP1, again demonstrating that the E2-TopBP1 interaction regulates E2 stability.

## Results

### Development of functional assays for investigating E2 segregation function

Previously we demonstrated co-localization of E2 and TopBP1 on mitotic chromatin, and shown that E2 can recruit TopBP1 to mitotic chromatin (41, 42). There were no assays available to determine whether E2 recruits plasmids onto mitotic chromatin; we therefore developed an assay. Using Label IT Tracker (Mirus, cat. no. MIR7025) we covalently added a fluorescent tag to ptk6E2-luc, a plasmid containing 6 E2 DNA binding sites upstream from the luciferase gene controlled by the tk promoter (43). We transfected this labeled plasmid into U2OS-Vec (vector control), U2OS-E2-WT (stably expressing wild type E2) and U2OS-E2-S23A (stably expressing E2 with serine 23 mutated to alanine, disrupting interaction with TopBP1) cell lines. Three days later we fixed the transfected cells and looked for the presence of the fluorescent ptk6E2-luc (Figure 1A). There was notable nuclear presence of the labeled plasmid in U2OS-E2-WT and U2OS-E2-S23A, but none in the U2OS-Vec cells. This suggests that both E2WT and E2S23A can bind to the labeled plasmid and retain it in the nucleus. Both E2WT and E2S23A activate transcription to an identical level in a 3-day transcription assay in U2OS cells (42). However, it was noticeable that in U2OS-E2-WT mitotic cells, but not in U2OS-E2-S23A mitotic cells, the fluorescent ptk6E2-luc was retained on the mitotic chromatin (mitotic cells indicated with white arrows). To investigate this further, confocal images were taken at a higher resolution (Figure 1B). As expected, no fluorescence was detected on mitotic U2OS-Vec cells (top panels). However, in U2OS-E2-WT cells there was a clear retention of the fluorescent plasmid on the mitotic chromatin (middle panels), while in U2OS-E2-S23A cells this phenotype was absent (bottom panels). Retention of fluorescent ptk6E2-luc was detectable on mitotic chromatin in repeated U2OS-E2-WT cells, but never in U2OS-E2-S23A cells. To confirm that E2-WT was not promoting integration of the fluorescent plasmid into the host genome, we washed the fixed cells with 1M salt. As this wash breaks non-covalent interactions and removes the fluorescent DNA from the chromatin of U2OS-E2-WT cells, we conclude that the plasmid has not become integrated (Figure 1C). Finally, we labeled pHPV16LCR-luc (a plasmid containing the luciferase gene under the transcriptional control of the HPV16 long control region (42)) as previously with ptk6E2-luc and transfected the labeled plasmid into U2OS-Vec, U2OS-E2-WT and U2OS-E2-S23A (Figure 1D). Similar to ptk6E2-luc, pHPV16LCR-luc was retained in the nuclei by E2-WT and E2-S23A but was only retained on mitotic chromatin in E2-WT expressing cells. Again, this was reproducible in multiple mitotic cells; E2-WT retained the fluorescent plasmid on chromatin while E2-S23A could not. In these experiments, we could not co-stain for E2 protein as the permeabilization disrupted the interaction of the fluorescent plasmids with the chromatin. These cell lines stably express E2, and most cells retain E2 protein expression (42).

**Figure 1.**
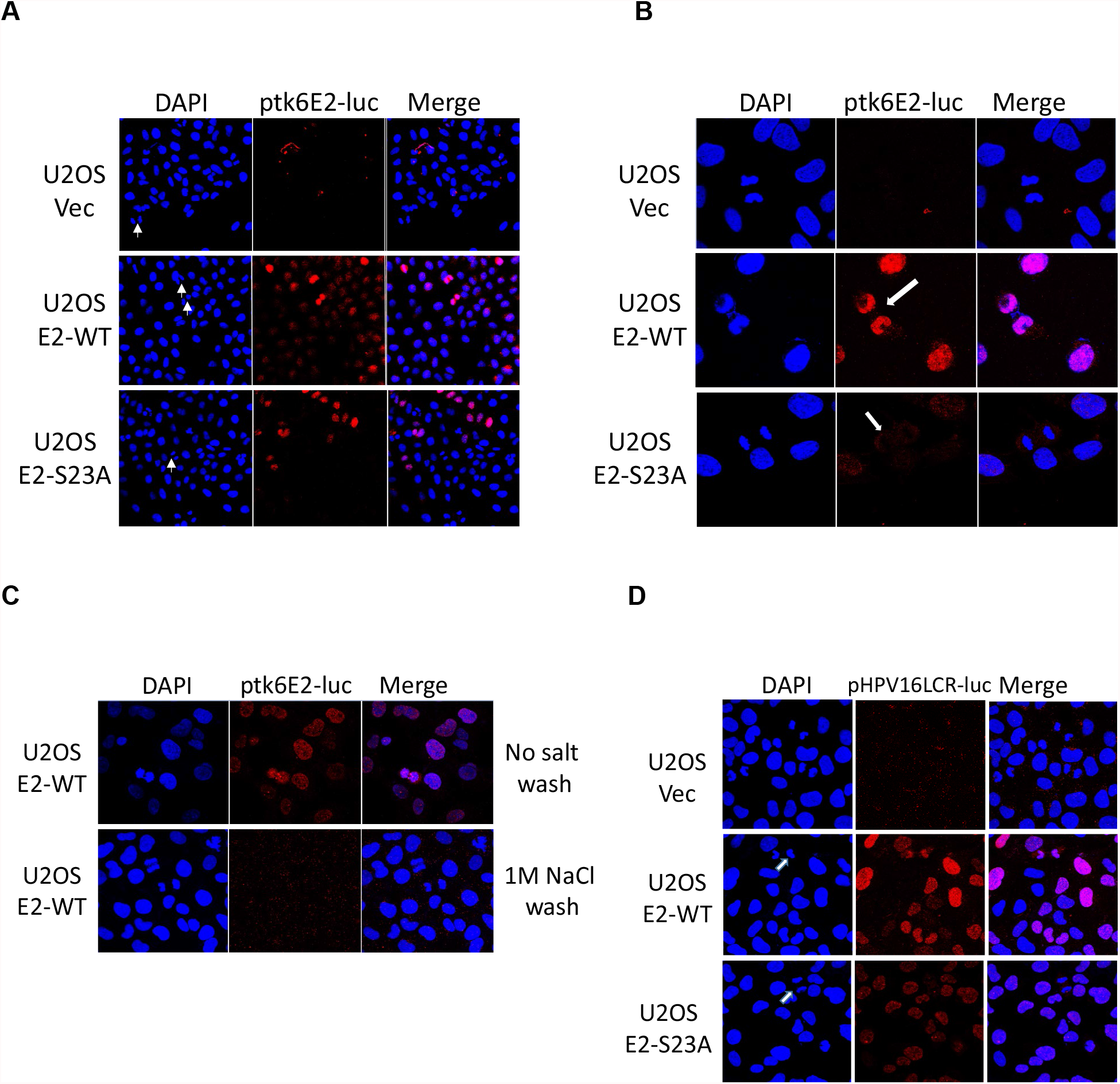
E2 recruits E2 binding site plasmids to U2OS cell mitotic chromatin in a TopBP1 interacting dependent manner. A. Fluorescently labeled ptk6E2-luc was transfected into the indicated cell lines. Three days later, Keyence imaging system (BZ-X810) captured the images shown. Mitotic cells are indicated with white arrowheads in the left panel. B. Confocal images demonstrate recruitment of ptk6E2-luc to mitotic chromatin in U2OS-E2-WT cells (arrow in middle panel). This never occurred in U2OS-E2-S23A cells. C. Washing with 1M NaCl removes fluorescent ptk6E2-luc from chromatin in U2OS-E2-WT cells demonstrating the plasmid is not integrated. D. Fluorescently labeled pHPV16LCR-luc is recruited to mitotic chromatin in U2OS-E2-WT cells but not in U2OS-E2-S23A (white arrows in left panels highlights mitotic cells).

When there is no selective pressure present, transfected plasmids are quickly lost from cells after 3-4 days (44). Due to the retention of ptk6E2-luc on mitotic chromatin by E2-WT (Figure 1), we investigated whether this led to a retention of the plasmid beyond three days when compared with U2OS-Vec and U2OS-E2-S23A. Figure S1A demonstrates that 6 days following transfection there remained a significant number of fluorescent plasmid positive cells in U2OS-E2-WT cells, but none in the U2OS-Vec or U2OS-E2-S23A, while Figure S1B demonstrates retention of fluorescent ptk6E2-luc by E2-WT on mitotic chromatin. Nine days following transfection there was a marked reduction, but still detectable, level of fluorescent ptk6E2-luc in the E2-WT cells (Figure S1C), and the fluorescent DNA could still be detected on mitotic DNA in the E2-WT cells (Figure S1D). Finally, we fluorescently labeled pSV40-luc (pGL3-Control from Promega, containing the SV40 promoter and enhancer), a plasmid that does not contain E2 DNA binding sites, and transfected it into U2OS-E2-WT. This plasmid did not associate with mitotic chromatin in E2-WT cells at day 3 (Figure S1E). It occurred to us that this retention of ptk6E2-luc by E2-WT would allow us to quantitate retention by measuring luciferase activity. On this principle, we developed a novel quantitative luciferase-based assay to measure the plasmid segregation/retention function of E2-WT (Figure 2A). In this assay, ptk6E2-luc or pSV40-luc are transfected independently into either U2OS-E2-WT or U2OS-E2-S23A. We could not use U2OS-Vec cells in this assay, as the luciferase activity of ptk6E2-luc is very low in the absence of E2, which made the measurement of plasmid retention via luciferase detection impractical. Three days following transfection, the cells were trypsinized and half of them used for a luciferase assay, and half of them replated. At day 6, cells were trypsinized and half of them used for a luciferase assay, and half of them replated. At day 9, all cells were harvested for luciferase assays. The ability of E2-WT and E2-S23A to activate transcription over a day three period is identical, this was therefore set as the reference point 1, to which the activity at day 6 and 9 is presented as relative (42). The upper panel of Figure 2A demonstrates that, in U2OS-E2-WT cells, around 85% of luciferase activity is retained between days 3 and 6, while in U2OS-E2-S23A cells retention is around 15% (note the log scale). At day 9 there remained around a 15% retention of luciferase activity in E2-WT cells, while E2-S23A cells had lost almost all activity when compared with day 3 levels. The lower panel demonstrates that for pSV40-luc 90% of activity is lost between days 3 and 6 irrespective of E2 type, and almost all activity is lost by day 9. These results reflect the retention of ptk6E2-luc by E2-WT, as indicated by the fluorescence assays described in Figures 1 and S1.

**Figure 2.**
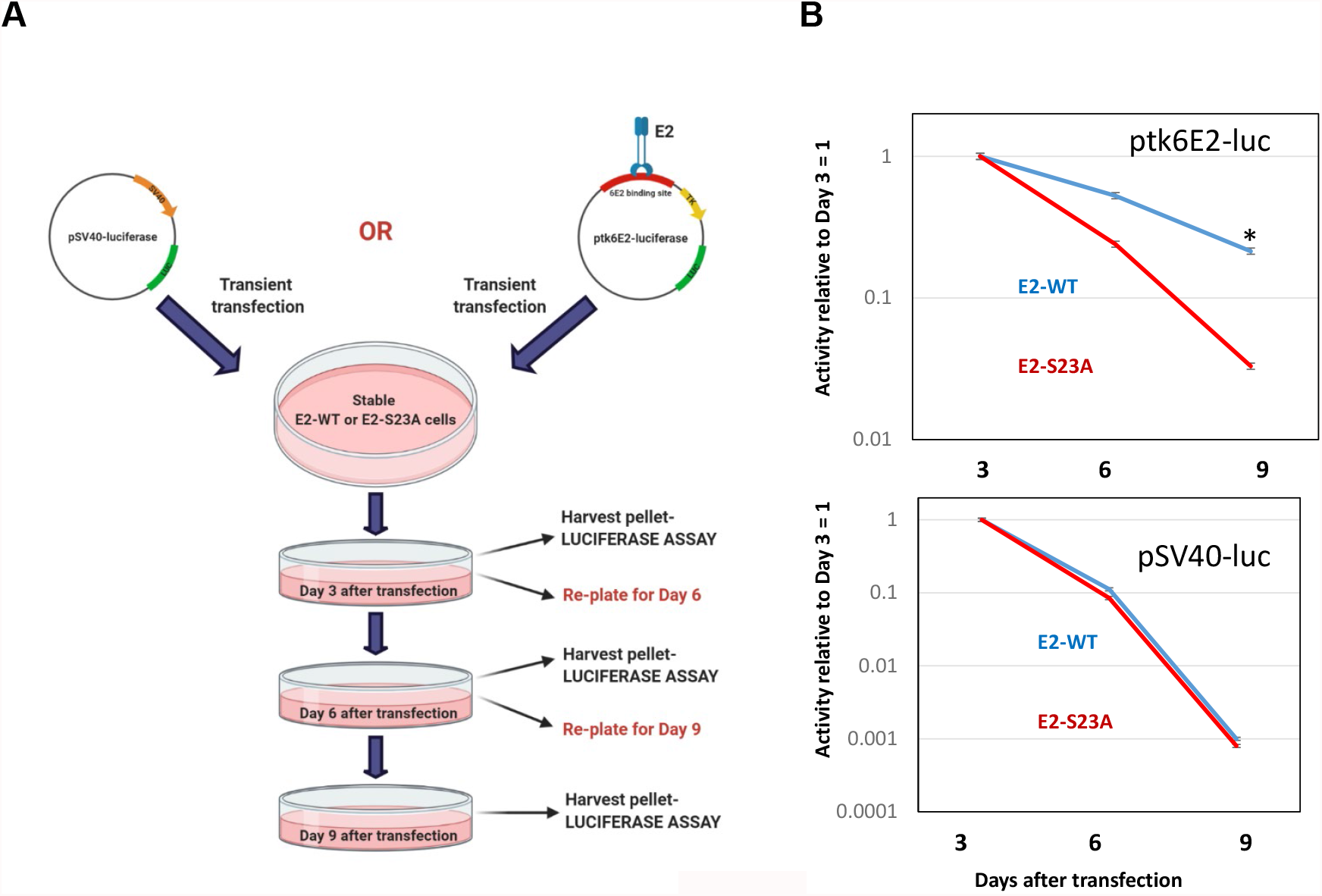
Plasmid retention/segregation by E2 in U2OS cells. A. This figure summarizes our luciferase-based segregation/retention assay summarized in the text. B. Luciferase activity detected in U2OS cells. The day 3 luciferase activity for both E2-WT and E2-S23A was statistically identical (42). For ptk6E2-luc, at day 6, E2-WT retained around 85% of day 3 activity while E2-S23A retained around 15%. At day 9, E2-WT retained around 15% of day 3 activity while most activity was lost with E2-S23A. For pSV40-luc, activity was lost rapidly in both E2-WT and E2-S23A cells. * indicates a significant difference between this sample and the others at days 6 and 9, p-value<0.05. The standard error bars are too small to show up on the log scale, the experiments represent the summary of three independent experiments.

### TopBP1 interaction is required for the plasmid retention/segregation function of E2

We set out to confirm that interaction with TopBP1 is essential for the retention/segregation function of E2-WT using our luciferase assay. In order to do this, we knocked out TopBP1 using siRNA. Figure 3A confirms that, following TopBP1 knockdown, E2 is removed from mitotic chromatin; this was observed in multiple mitotic cells. We next carried out the assay described in Figure 2A, but introduced TopBP1 or scrambled control siRNA to the cells at the day 3 time-point prior to replating. Figure 3B demonstrates that at day 6 following transfection (3 days following addition of siRNA) there was a significant knock down of TopBP1 (compare lanes 4-6, TopBP1 siRNA, with 1-3, scr siRNA). This knockdown extended to day 9 (lanes 7-9). Figure 3C (upper panel) demonstrates that knockdown of TopBP1 expression abrogated the retention of ptk6E2-luc luciferase activity when compared with the scr siRNA control. Knockdown of TopBP1 made no difference to the activity detected with E2-S23A at either time point. Similarly, knockdown of TopBP1 had no effect on the activity of pSV40-luc in either of the E2 expressing lines at either time point. During the time course of the TopBP1 siRNA knockdown there was no significant alteration of cellular growth due to the absence of TopBP1 (Figure S2A). We repeated the TopBP1 knockdown with two additional TopBP1 targeting siRNAs; these experiments produced identical results to those in Figure 3C (Figure S2B).

**Figure 3.**
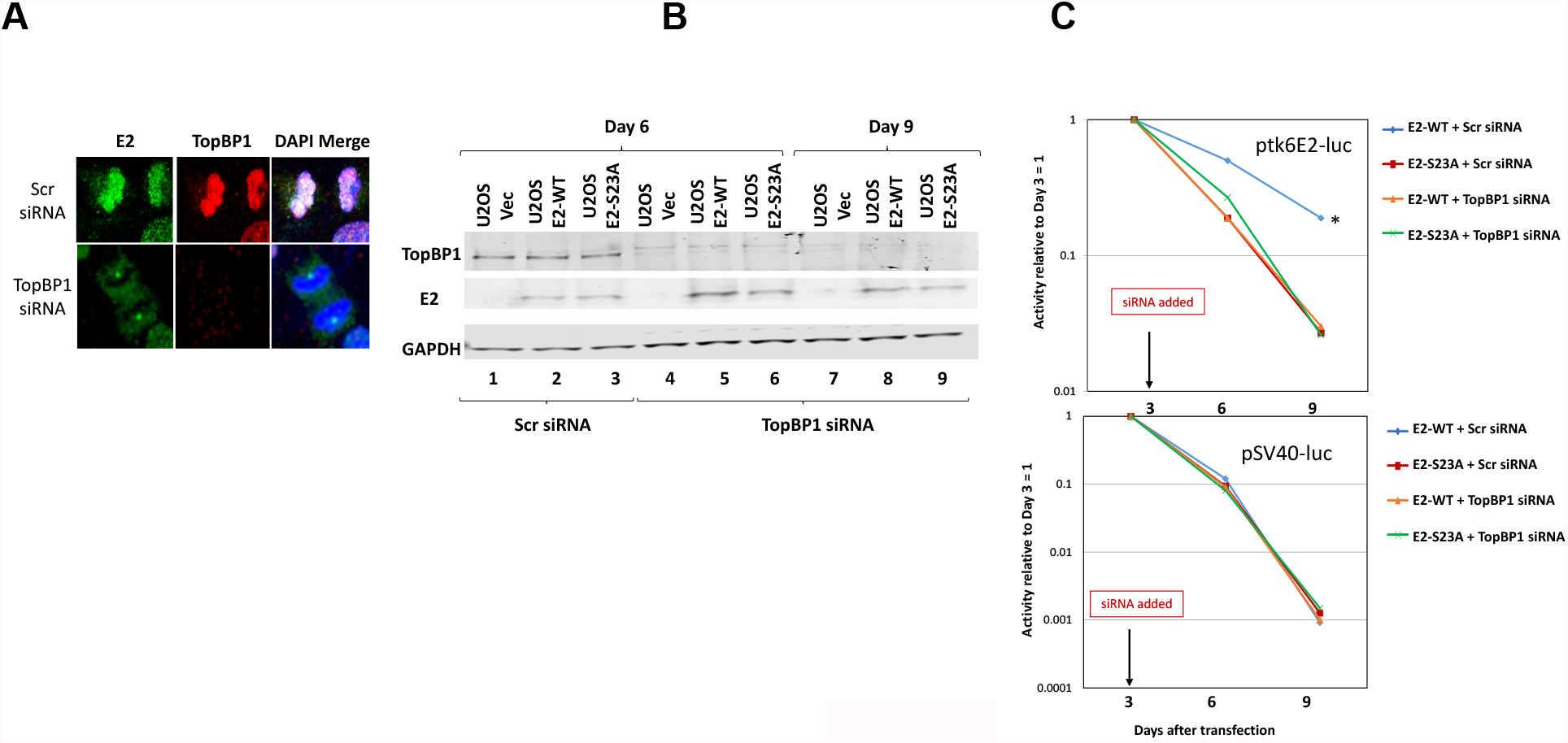
E2 plasmid segregation/retention function is dependent upon TopBP1 in U2OS cells. A. siRNA knockdown of TopBP1 (bottom panels) removes the interaction of E2-WT with mitotic chromatin. B. siRNA was added to cells at day-3 of our luciferase-based plasmid retention/segregation assay described in Figure 2A. Protein was then prepared from cells 6 days and 9 days following transfection and western blotting for the indicated proteins carried out. The TopBP1 siRNA knocked down TopBP1 expression that persisted from day 6 until day 9. C. The knockdown of TopBP1 expression abolishes the ability of E2-WT to retain ptk6E2-luc. * indicates a significant difference between this sample and the others at days 6 and 9, p-value<0.05. The standard error bars are too small to show up on the log scale, the experiments represent the summary of three independent experiments.

To further confirm that the E2-TopBP1 interaction is require for the E2-WT retention/segregation function we knocked down components of CK2, a kinase we demonstrated phosphorylates serine 23 of E2-WT to promote complex formation with TopBP1 *in vitro* and *in vivo* (42). CK2 functions as a tetramer that has two active catalytic components; *α* and *α*’ (45). It is not possible to knockout both *α* sub-units simultaneously as cells become unviable, therefore we knocked out both components individually and measured their effect on E2-WT plasmid retention/segregation function. Knockdown of either component compromises the E2-TopBP1 interaction as it reduces E2 phosphorylation on serine 23 (42). Figures 4A and 4B demonstrate knockdown of CK2a and CK2a’, respectively, during the time of the assay without significant disruption to E2 expression, similarly to TopBP1 (Figure 3B). Figure 4C demonstrates that knock-down of CK2*α* or *α*’ results in a compromised segregation/retention E2 function. Noticeably, at day 6 there was a total elimination of E2 retention function that somewhat recovers by day 9. This is perhaps due to the reorganization of the CK2 enzyme in the cells, arranging into tetramers with only CK2*α* or *α*’ components compensating for the loss of the other *α* component.

**Figure 4.**
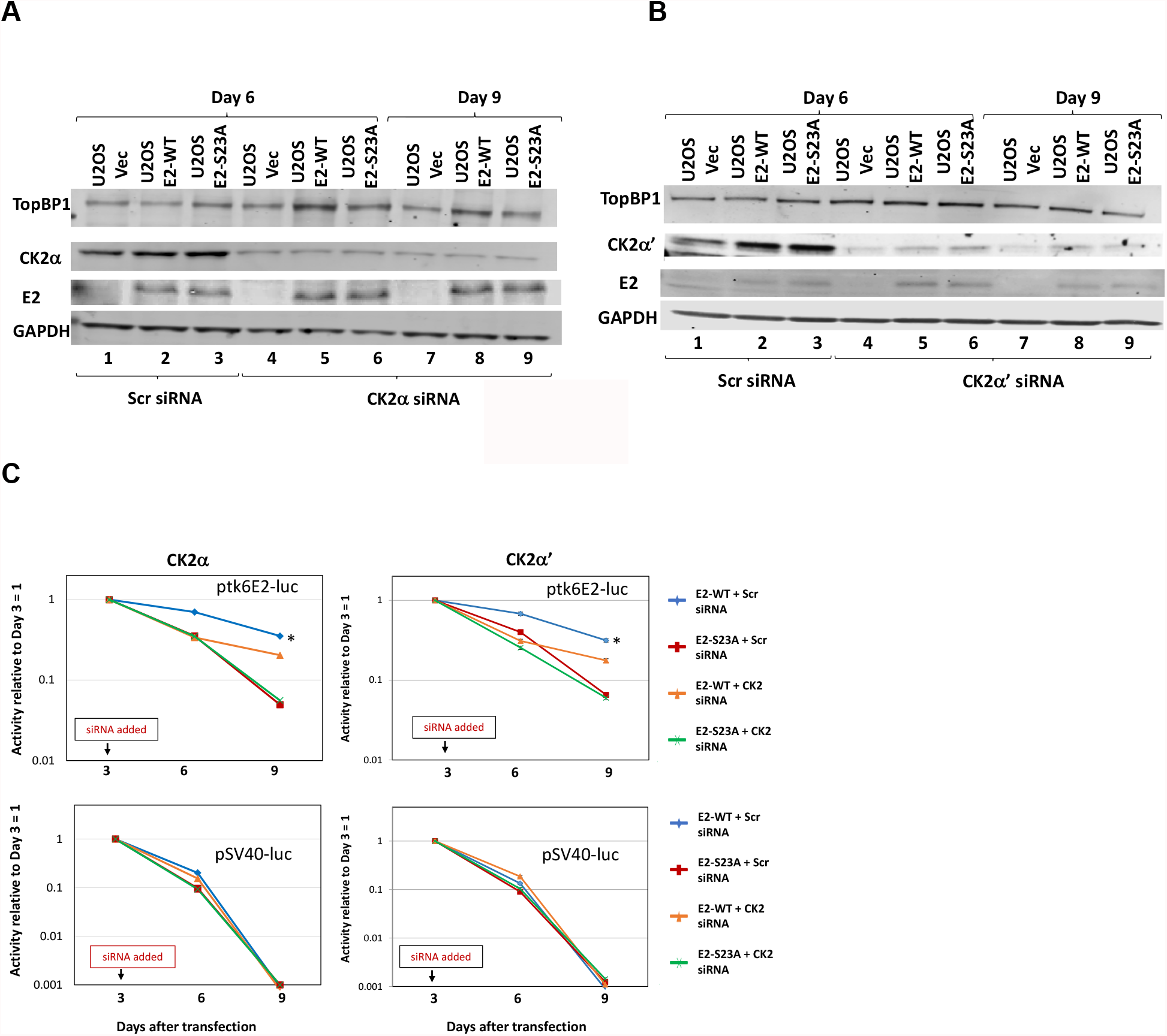
Knockdown of CK2 components disrupts E2 segregation/retention function in U2OS cells. A and B. Indicated siRNA was added to cells at day-3 of our luciferase-based plasmid retention/segregation assay described in Figure 2A. Protein was then prepared from cells 6 days and 9 days following transfection and western blotting for the indicated proteins carried out. C. The knockdown of CK2 components compromises the ability of E2-WT to retain ptk6E2-luc. * indicates a significant difference between this sample and the others at days 6 and 9, p-value<0.05. The standard error bars are too small to show up on the log scale, the experiments represent the summary of three independent experiments.

### The E2-TopBP1 interaction is critical for E2 segregation/retention function in human keratinocytes

Using N/Tert-1 cells stably expressing E2-WT or E2-S23A (42), we carried out our segregation/retention assay. Figure 5A demonstrates that in N/tert-1 cells E2-WT is able to retain ptk6E2-luc activity when compared with that in E2-S23A cells, similarly to U2OS cells (Figure 2). E2-WT protein expression levels in U2OS increase during mitosis, and TopBP1 levels also increase during mitosis in the presence of E2-WT. Neither E2 nor TopBP1 levels increase in E2-S23A cells. We identified a similar increase in E2 and TopBP1 in N/Tert-1-E2-WT cells 16 hours following release from a double thymidine block, neither protein increased in the E2-S23A cells (Figure 5B). Next, we wanted to determine whether this increase in E2 protein expression during mitosis occurs in cells immortalized by the HPV16 genome. Figure 6A demonstrates that this is the case. Human foreskin keratinocytes immortalized with HPV16 genomes (HFK+HPV16) were double thymidine blocked (DTB) and released for the indicated time points; this line has been described previously (42). Preliminary studies to help identify mitosis indicated that there was no change in E2 or TopBP1 expression levels before a 16-hour time point following release from the block (not shown). In Figure 6A, Cyclin B1 and pH3 (ser10) antibodies confirm that the increase in E2 and TopBP1 levels were occurring during mitosis. Another notable feature in this Figure is that the presence of 3T3-J2 fibroblasts increases the expression levels of the E2 protein (compare lanes 2 and 3). This suggests that the “stroma” used during organotypic rafting studies (collagen plugs infused with 3T3-J2 fibroblasts) cross talks with the keratinocytes to regulate the expression levels of E2. Our previous work demonstrated that, using immunostaining, there was no detectable E2 expression in HFK+HPV16-S23A cells (immortalized with a HPV16 genome containing the serine 23 to alanine mutation in E2) (42). We confirmed the lack of E2 expression by western blotting of HFK+HPV16 and HFK+HPV16-S23A cells double thymidine blocked and released into mitosis in monolayer cells grown with 3T3-J2 fibroblasts (Figure 6B). Strikingly, there was no E2 expression in HFK+HPV16-S23A cells. To confirm that this occurred during the viral life cycle, we extracted proteins from organotypic raft cultures and demonstrated no E2 expression in HFK+HPV16-S23A cells when compared with HFK+HPV16 (Figure 6C). We also confirmed that this loss of E2 expression substantially abrogates viral replication, as *γ*H2AX expression is lost in the HFK+HPV16-S23A cells. *γ*H2AX is a marker of active HPV16 replication. The E2 and *γ*H2AX blots were run separately resulting in the two GAPDH blots. While E2 expression was lost in the HFK+HPV16-S23A cells, viral genomes could still be detected by fluorescent in situ hybridization (42). To determine whether these genomes were integrated due to the loss of E2 expression and failure to replicate, we carried out Southern blots on DNA extracted from day 7 and day 14 rafts (Figure 6D). At day 7 rafts are not fully differentiated, while at day 14 they are, and viral genome amplification should have occurred (46). Figure 6D shows the outcomes from studies on two rafts generated from independently immortalized HFK cells. For HFK+HPV16-WT 1 and 2 there was an increase in the 8kbp band detected between days 7 and 14 demonstrating amplification of an episomal viral genome (compare lanes 1 with 5 and 3 with 7, respectively). Amplification was more with sample 2, but amplification occurred with both samples. However, for HFK+HPV16-S23A 1 and 2 the viral DNA was mostly integrated demonstrating a loss of the band at 8kbp. There is an 8kbp band detected in sample 1 after 14 days indicating there may be some episomal genomes that are amplified. Ultimately, the DNA is predominantly integrated in both samples. This is a striking observation for several reasons. In monolayer cultures of these lines, grown without 3T3-J2 fibroblasts, the viral genomes are episomal in all of these lines and there are, if anything, more viral genomes in the S23A samples. Please see (42) for details. Therefore, as soon as the cells are exposed to the collagen-fibroblast “stroma” required for organotypic rafting, E2 expression is lost and the viral genome begins to integrate. In HFK+HPV16-WT cells, addition of 3T3-J2 fibroblasts increases E2 protein levels to detectable levels (Figure 6A, compare lanes 2 and 3). Therefore, the stroma regulates E2 protein levels in HFK contributing to the control of the viral life cycle. Loss of E2 interaction with TopBP1 clearly results in a loss of E2 expression; the E2-TopBP1 interaction is critical for the HPV16 life cycle. It is noticeable that, in organotypic rafts of the HFK+HPV16-S23A cells, there is a more transformed epithelium perhaps driven by the integration of the viral genome; again, please see (42) for details.

**Figure 5.**
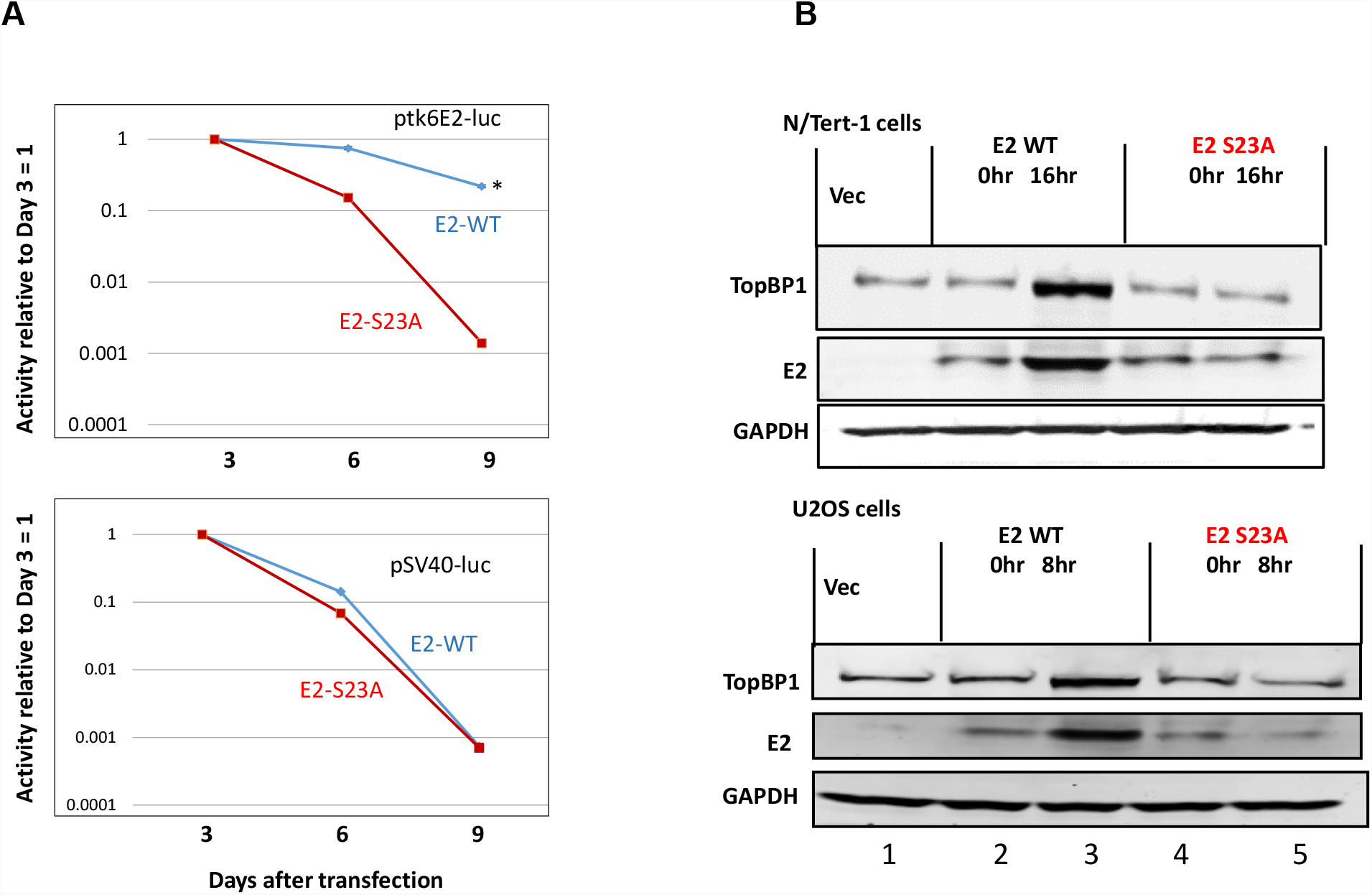
TopBP1 interaction regulates E2 plasmid segregation/retention function and mitotic expression in N/Tert-1 cells. A. Luciferase activity detected in N/Tert-1 cells. The day 3 luciferase activity for both E2-WT and E2-S23A was statistically identical (42). For ptk6E2-luc, at day 6, E2-WT retained around 90% of day 3 activity while E2-S23A retained around 15%. At day 9, E2-WT retained around 20% of day 3 activity while most activity was lost with E2-S23A. For pSV40-luc, activity was lost rapidly in both E2-WT and E2-S23A cells. B. N/Tert-1 (top panel) or U2OS (lower panel) were double thymidine blocked and released for the indicated time points where cells have progressed into mitosis (42). Harvested proteins were then western blotted for the indicated proteins and the results demonstrate an increase in E2-WT, but not E2-S23A, protein expression in mitosis. In A, * indicates a significant difference between this sample and the others at days 6 and 9, p-value<0.05. The standard error bars are too small to show up on the log scale, the experiments represent the summary of three independent experiments.

**Figure 6.**
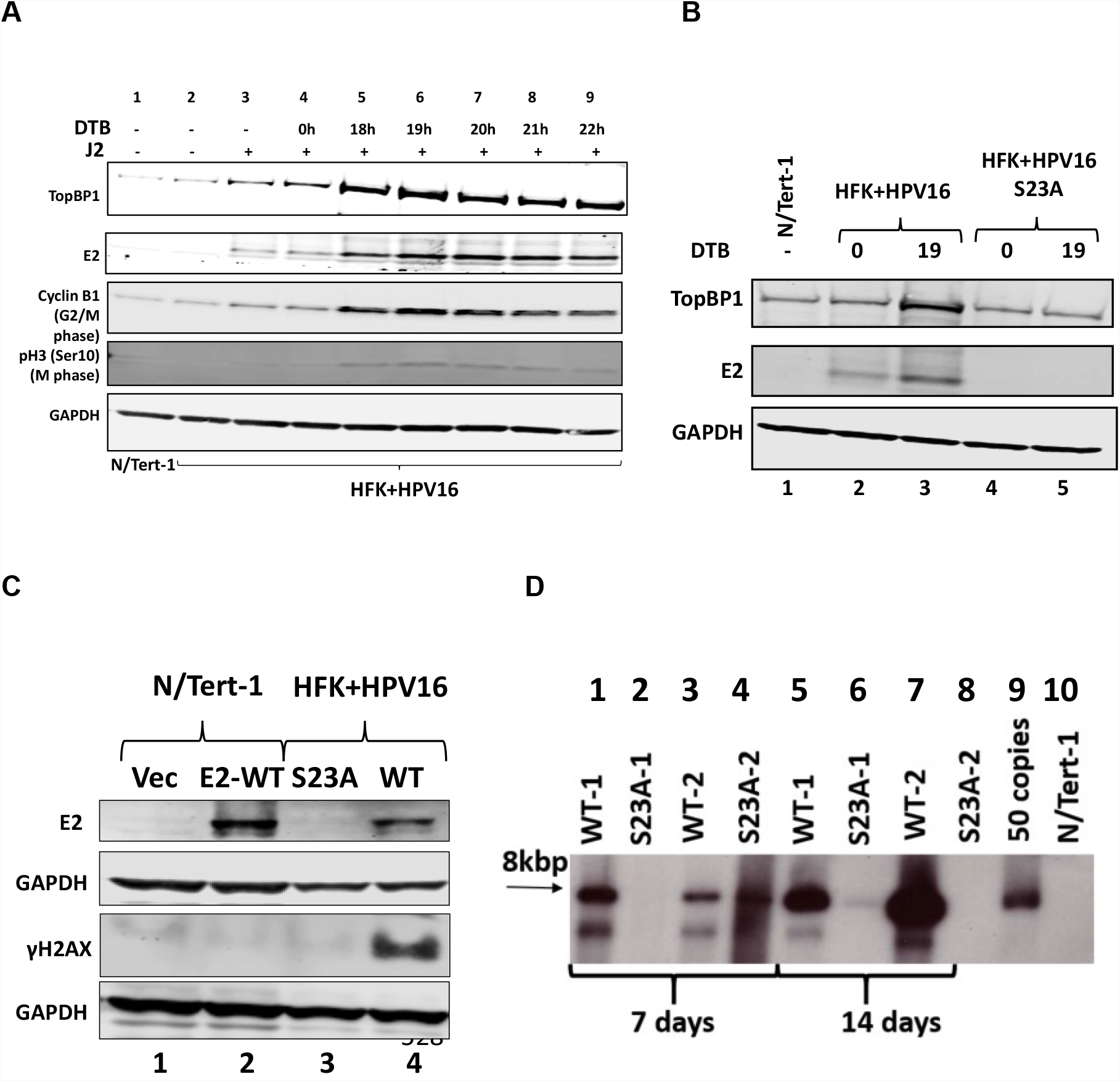
Interaction with TopBP1 regulates E2 stability during the cell cycle and the viral life cycle. A. Human foreskin keratinocytes immortalized by HPV16 (HFK+HPV16) (42) were double thymidine blocked and released for the indicated time points. Protein extracts were prepared and the indicated western blots carried out. Cyclin B and pH3 (Ser10) indicate G2/M and M, respectively. B. HFK+HPV16 and HPV16+HPV16-S23A (E2 serine 23 mutated to alanine) grown with J2 fibroblasts were double thymidine blocked and released for 19 hours (mitosis) and blotted with the indicated proteins. 0 = double thymidine blocked, 19 = 19 hours following release from block. C. N/Tert-1 Vector control cells and E2-WT expressing cells were used as an E2 detection control (lanes 1 and 2). Lane 3 is a blot of proteins extracted from an organotypic raft of HFK+HPV16-S23A immortalized cells, while lane 4 is an extract from an HFK+HPV16 organotypic raft. The extracted proteins were blotted with the indicated antibodies. D. DNA was harvested from the indicated samples at day 7 (partial differentiation and life cycle) and day 14 (full differentiation and life cycle) and Southern blotting carried out using an entire HPV16 genome.

We next investigated whether it was the addition of 3T3-J2 fibroblasts or induction of differentiation in the organotypic rafts that induces E2 degradation. To do this, N/Tert-1+E2-WT and N/Tert-1+E2-S23A cells were calcium differentiated in monolayer with or without 3T3-J2 fibroblasts. Figure 7A demonstrates that, even in cells that have not undergone differentiation, there is a loss of E2-S23A protein following addition of 3T3-J2 cells (compare lanes 3 and 6). Involucrin was used as a loading control in these experiments due to the differentiation experiments. Addition of MG132 to the N/Tert-1+E2-S23A cells restored E2 expression demonstrating that the addition of 3T3-J2 fibroblasts promoted E2 degradation via the proteasome (Figure 7B). To confirm that 3T3-J2 degradation of E2-S23A was due to a compromised interaction with TopBP1, we used siRNA to downregulate TopBP1 in N/Tert-1+E2-WT cells and added 3T3-J2 fibroblasts. Figure 7C demonstrates that downregulation of TopBP1 reduces E2 expression levels following the addition of 3T3-J2 fibroblasts (compare lane 2 with 3).

**Figure 7.**
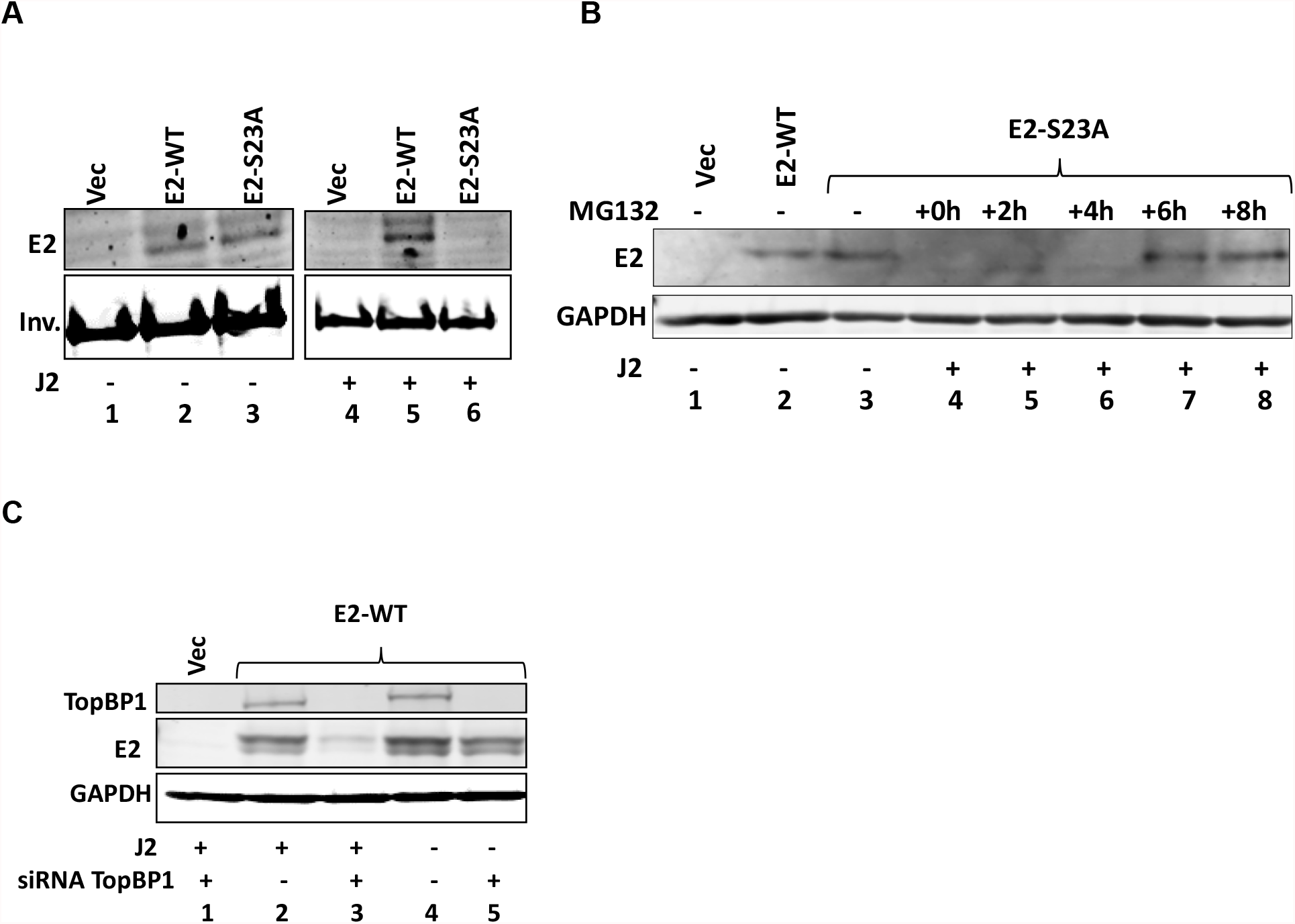
Interaction with TopBP1 regulates E2 proteasomal turn over in the presence of J2 fibroblasts. A. N/Tert-1 Vector control (Vec), E2-WT or E2-S23A cells were grown in the absence (lanes 1-3) and presence (lanes 4-6) of J2 fibroblasts. Protein extracts were prepared and blotted with the indicated antibodies. B. MG132 addition restores expression of E2-S23A in N/Tert-1 cells in the presence of J2 fibroblasts. Addition of J2 cells eliminates detectable E2-S23A expression (compare lanes 3 and 4), and E2 expression is recoverable after 6 hours of MG132 treatment (lane 7). C. TopBP1 siRNA reduces TopBP1 protein expression (lanes 3 and 5), and the addition of J2 fibroblasts in the absence of TopBP1 expression reduces E2-WT protein levels (compare lanes 3 and 5.

## Discussion

While E2 interaction with mitotic chromatin suggests it is a plasmid segregation/retention factor during the HPV16 life cycle (41), to date there is little evidence that E2 has such a function. To our knowledge, there is only one manuscript describing assays for measuring E2 segregation/retention function, and that was for BPV1 E2 (47). In this assay, GFP expression was monitored in a plasmid that contained the GFP gene, E2 DNA binding sites, and an E2 expression cassette. This study, with BPV1 E2, indicated there were two functions of E2 contributing to the plasmid segregation/retention function of BPV1 E2, interaction with mitotic chromatin by itself was insufficient. We are able to generate stable cells expressing E2-WT and mutant proteins enabling us to dissociate E2 expression from any reporter plasmids used in segregation/retention assays (18, 42, 48-50). We exploited these cell lines to demonstrate that E2 protein has plasmid segregation/retention function and that this activity was dependent upon E2 interaction with TopBP1. Figure 1 demonstrates that E2 binding site containing fluorescent plasmids are recruited to mitotic chromatin by E2-WT, but not by the TopBP1 interacting deficient E2-S23A. Figure S1 demonstrates that E2-WT retains fluorescent signals even 9 days following the original transfection. Figure 2 describes exploiting the ptk6E2-luc plasmid to develop a quantitative assay for measuring E2 segregation/retention function, and this again demonstrates that E2-S23A has lost this function. To confirm that the failure of E2-S23A in these assays is due to a failure to interact with TopBP1 we took two approaches. First, in Figure 3 we used siRNA to knockdown TopBP1 expression, resulting in a loss of the segregation/retention function of E2-WT. We repeated this with two additional TopBP1 siRNAs and generated identical results (Figure S2). Second, Figure 4 demonstrated that siRNA knock down of CK2 components disrupted E2-WT segregation/retention function. Previously, we demonstrated that CK2 phosphorylation of E2 on serine 23 is required for E2-TopBP1 complex formation. Overall, these results demonstrate that E2-WT has segregation/retention function mediated via interaction with TopBP1.

All of the results discussed above were carried out in U2OS cells, we next moved this project into human keratinocytes, the natural target cell for HPV16 infection and disease. Previously, we demonstrated that in N/Tert-1 cells (human foreskin keratinocytes immortalized with hTERT) E2-WT interacted with TopBP1, while E2-S23A could not (42). We also demonstrated that CK2 phosphorylated E2 on serine 23 in N/Tert-1 cells (42). Figure 5 demonstrates, using our luciferase-based plasmid segregation/retention assay, that E2-S23A has lost the ability to mediate this function, just as it has in U2OS cells. We previously reported that E2 and TopBP1 protein levels increased during mitosis in U2OS cells in an interaction dependent manner, and we demonstrate that in N/Tert-1 cells this is also true (Figure 5). This increase in E2 and TopBP1 during mitosis also occurs in HFK+HPV16 cells (Figure 6). There were also several striking observations in N/Tert-1 and HFK+HPV16 cells with regard interaction with stromal 3T3-J2 fibroblasts used in organotypic rafting experiments. In HFK+HPV16-S23A cells grown in monolayer with 3T3-J2 fibroblasts, E2 expression is lost (Figure 6). E2 expression is also lost in organotypic rafting of HFK+HPV16-S23A demonstrating that the E2-TopBP1 interaction is critical for the viral life cycle. We demonstrate that this loss of E2-S23A expression is due to a failure to interact with TopBP1, as TopBP1 siRNA knockdown results in a reduction of E2-WT expression in N/Tert-1 cells in the presence of 3T3-J2 fibroblasts. The loss of E2-S23A expression in N/Tert-1 cells co-cultured with 3T3-J2 fibroblasts is due to enhanced proteasomal turnover of the protein, as the addition of MG132 rescues E2-S23A expression in the N/Tert-1 – 3T3-J2 co-cultured cells. The knockdown of E2 expression due to a failure of interaction with TopBP1 has a catastrophic effect on the viral life cycle, as viral replication is abrogated and the viral genomes integrate into that of the host during organotypic rafting. This integration likely contributes to the transformed phenotype of HFK+HPV16-S23A organotypic raft cultures (42).

Given that E2-S23A is degraded during the viral life cycle, it is impossible to determine whether the segregation/retention function of E2 is critical during the viral life cycle using this mutant. Our results demonstrate that E2 encodes such a function, but further studies are required to determine the role of this function during the viral life cycle. Any disruption of the E2-TopBP1 interaction would result in a reduction in E2 protein expression. We are currently working on identifying the lysine residues responsible for mediating the proteasomal degradation of E2-S23A and will incorporate a mutation of these lysines into the S23A background to generate an E2 protein that fails to interact with TopBP1, but is expressed during the viral life cycle. The increased expression of E2 during mitosis is also under further investigation, the increase in expression is reminiscent of enhanced acetylation that we have observed following knockout of SIRT1 expression in E2 expressing cells (51). This stabilization of E2 during mitosis does not occur with E2-S23A, it is possible that the mechanisms that regulate E2 stability during mitosis and following addition of 3T3-J2 fibroblasts to keratinocytes are linked, and we are currently investigating this.

Overall, the novel assays described here confirm that E2 has plasmid segregation/retention activity and that an interaction with TopBP1 is critical for this function. There are likely other factors involved in the E2-TopBP1 mitotic complex that are involved in mediating the segregation/retention function. For example, we recently demonstrated that TopBP1 and BRD4 form a complex and are currently investigating the role of this interaction in the plasmid segregation function of E2 (42).

## Materials and methods

### Cell culture

Stable cell lines expressing wild type E2 (E2-WT) and E2-S23A (E2 with serine 23 mutated to alanine, abrogating interaction with TopBP1), alongside cells with pcDNA empty vector plasmid control were established both in U2OS and N/Tert-1 cell lines as previously described (18, 42). Cell culture was as described in these publications. For immortalization and culture of human foreskin keratinocytes with HPV16 E2-WT and HPV16 E2-S23A, see (42).

### Generation of fluorescently tagged plasmids and transfection

Label IT Tracker (Mirus, cat. no. MIR7025) protocol was used to covalently attach a fluorescein-492 tag to ptk6E2-luc, a plasmid containing 6 E2 DNA binding sites upstream from the luciferase gene controlled by the tk promoter (43). This fluorescent ptk6E2-luc plasmid was then transiently transfected into following stable cells; U2OS-Vec (vector control), U2OS-E2-WT (stably expressing wild type E2) and U2OS-E2-S23A. After 48h of transfection, the transfected cells were passaged for next timepoint and a separate set of cells grown on coverslips were simultaneously fixed, stained with DAPI and observed for the presence of the fluorescent ptk6E2-luc by immunofluorescence, The above procedure was followed to label pHPV16LCR-luc (a plasmid containing the luciferase gene under the transcriptional control of the HPV16 long control region (29) and transfected the labeled plasmid into U2OS-Vec, U2OS-E2-WT and U2OS-E2-S23A as mentioned above.

### Immunofluorescence

U2OS cells expressing stable E2-WT, E2-S23A and pcDNA empty vector plasmid control were plated on acid-washed, *poly-L-lysine* coated coverslips, in a 6-well plate at a density of 2 × 10^5^ cells / well (5 ml DMEM + 10% FBS media). After 48h of transfection, the cells were washed twice with PBS, fixed, and stained as described (42). The primary antibodies used are as follows: HPV16 E2 B9 monoclonal antibody, 1:500 (42), TopBP1 1:1000 (Bethyl; catalog no. A300-111A). The cells were washed and incubated with secondary antibodies Alexa fluor 488 goat anti-mouse (Thermo fisher; catalog no. A-11001) and Alexa fluor 594 goat anti-rabbit (Thermo fisher; catalog no. A-11037) diluted at 1: 1000. The wash step was repeated, and the coverslips were mounted on a glass slide using Vectashield mounting medium containing 4′,6-diamidino-2-phenylindole (DAPI). Images were captured with a Keyence imaging system (BZ-X810) or using Zeiss LSM700 laser scanning confocal microscope and analyzed using Zen LE software.

### Western blotting

Protein was harvested from cell pellets lyzed using 2x pellet volume NP-40 lysis buffer (0.5% Nonidet P-40, 50mM Tris [pH 7.8], and 150mM NaCl) supplemented with protease inhibitor (Roche Molecular Biochemicals) and phosphatase inhibitor cocktail (Sigma). The cells were lyzed for 20 min on ice followed by centrifugation at 18,000 rcf (relative centrifugal force) for 20 min at 4°C. Protein concentration was estimated colorimetrically using a Bio-Rad protein assay. 50 µg of protein with equal volume of 4X Laemmli sample buffer (Bio-Rad) was denatured at 95^°^C for 5 min. Proteins were separated using a Novex™ WedgeWell™ 4 to 12% Tris-glycine gel (Invitrogen) and transferred onto a nitrocellulose membrane (Bio-Rad) using the wet-blot method, at 30 V overnight. The membrane was blocked with *Li-Cor* Odyssey® blocking buffer (PBS) diluted 1:1 v/v with PBS and then incubated with specified primary antibody in *Li-Cor* Odyssey® blocking buffer (PBS) diluted 1:1 with PBS. Next, the membrane was washed with PBS supplemented with 0.1% Tween20 and probed with the Odyssey secondary antibodies (IRDye® 680RD Goat anti-Rabbit IgG (H + L), 0.1 mg or IRDye® 800CW Goat anti-Mouse IgG (H + L), 0.1 mg) in *Li-Cor* Odyssey® blocking buffer (PBS) diluted 1:1 with PBS at 1:10,000 for 1 h at room temperature. After washing with PBS-tween, the membrane was imaged using the Odyssey^®^ CLx Imaging System and ImageJ was used for quantification. Primary antibodies used for western blotting studies are as follows: monoclonal B9 1:500 (42), TopBP1 1:1000 (Bethyl; catalog no. A300-111A), GAPDH 1:10,000 (Santa Cruz; catalog no. sc-47724), Casein kinase IIα (1AD9) 1:500 (Santa Cruz; catalog no. sc-12738), CKII alpha’ antibody 1:1000 (Bethyl; catalog no. A300-199A), Histone H2A.X antibody (938CT5.1.1) 1:200 (Santa Cruz; catalog no. sc-517336), Involucrin antibody (SY5) 1:200 (Santa Cruz; catalog no. sc-21748), Cyclin B1 (D5C10) XP® Rabbit mAb 1:1000 (Cell Signaling Technology; catalog no. 4138) and phospho-histone H3 (Ser10) antibody 1:1000 (Cell Signaling Technology; catalog no.9701).

### Plasmid segregation assay

Two luciferase reporter plasmids were used for our novel assay; one containing the SV40 promoter and enhancer (pGL3 Control, Promega, described as pSV40-luc in this manuscript) which has no E2 DNA binding sites, the other with the HSV1 tk promoter driving expression of luciferase with 6-E2 target sites upstream; see results for details. The pSV40-luc and the ptk6E2-luc were transiently transfected, separately, into either E2-WT or E2-S23A cells. 3 days post-transfection, the cells were trypsinized and half re-plated and half harvested for a luciferase assay system (Promega). This luciferase activity was the baseline activity. At day 6, the same process was repeated, half of the cells harvested for luciferase assay, half re-plated. At day 9, cells were harvested for luciferase activity assay. Bio-Rad protein estimation assay was used for protein concentration estimation, to standardize for cell number. Relative fluorescence units were measured using the BioTek Synergy H1 hybrid reader. The activities shown are expressed relative to the respective protein concentrations of the samples. The assays shown are representative of three independent experiments carried out in triplicates.

### Small interfering RNA (siRNA) and segregation assay

U2OS parental cells were plated on a 100-mm plates. The next day, cells were transfected using 10 µM of siRNA in the table below. 10 µM of MISSION^®^ siRNA Universal Negative Control (Sigma-Aldrich; catalog no. SIC001) was used as a “non-targeting” control in our experiments. Lipofectamine™ RNAiMAX transfection (Invitrogen; catalog no. 13778-100) protocol was used in the siRNA knockdown. 48h post transfection, the cells were harvested, and knockdown confirmed by immunoblotting for the protein of interest. Segregation assays were performed as described before, after treating the cells with the siRNA of interest on day 3 of the protocol.

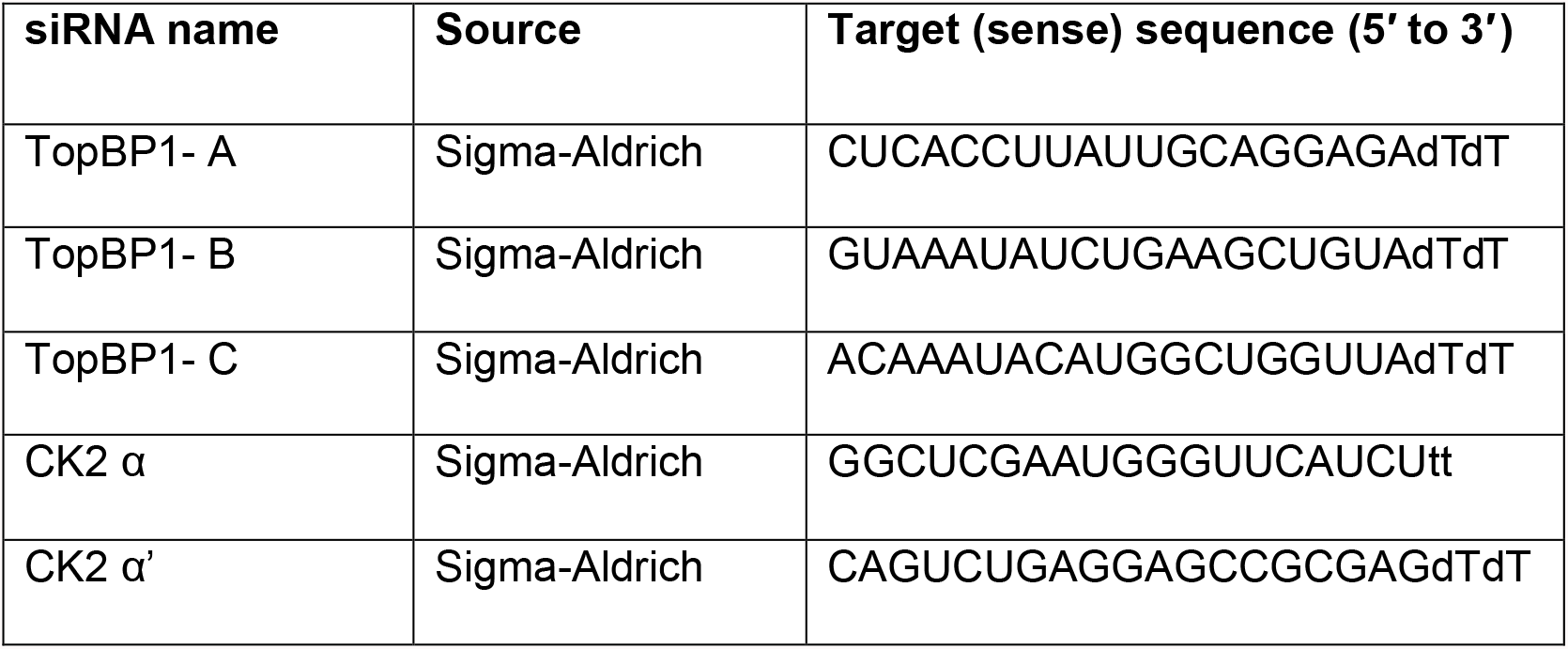

### Cell synchronization

U2OS and N/Tert-1 cells expressing stable E2-WT, E2-S23A and pcDNA empty vector plasmid control were plated at 3 × 10^5^ density onto 100-mm plates in DMEM + 10% FBS media. The cells were treated with 2 mM thymidine diluted in the supplemented DMEM media for 16 h. The cells were then washed 2 times with PBS and recovered in supplemented DMEM media. After 8 h, to block the cells at G1/S phase, a second dose of 2 mM thymidine was added and incubated for 17 h. The cells were then washed twice with PBS and recovered as before at the following time points: for U2OS, 0 h (G1/S phase) and 8 h (M1 phase). For N/Tert-1, cells were harvested at 0 h (G1/S phase) and 16 h (M1 phase). The cell lysates were prepared using the harvested cells at the time points mentioned and immunoblotting was carried out. The cell cycle phase analysis was carried out as previously described (CK2 paper). The above procedure was repeated in HFK cells immortalized with HPV16 genomes (HFK+HPV16 E2-WT or HFK+HPV16 E2-S23A) grown with or without 3T3-J2 fibroblasts. N/Tert-1 vector control cells were used as control. Cyclin B1 and pH3 (ser10) antibodies were used to confirm the mitosis phase in these cells using immunoblotting.

### MG132 proteasomal inhibitor treatment

N/Tert-1 cells expressing stable E2-S23A were plated on mitomycin C-treated J2 feeders in KSFM media. Next day, the cells were treated with 10μM of MG132 and harvested at 0h, 2h, 4h, 6h, and 8h post-treatment and immunoblotting was carried out to detect E2 and GAPDH.

### Organotypic raft culture

Keratinocytes were differentiated via organotypic raft culture as described previously (mSphere 2020). Briefly, one million cells were seeded onto collagen matrices containing one million3T3-J2-3T3 fibroblast feeder cells. Cells were then grown to confluency atop the collagen matrices, indicated by media colour change. Plugs were then lifted onto wire grids and cultured at the air-liquid interface for 13 days, with media replacement on alternate days. For protein extraction, the skin layer of cells that had grown was separated from the collagen plug and homogenized using a dounce homogenizer, in UTB buffer (9 M urea, 50 mM Tris, pH 7.5, 150 mM β-mercaptoethanol) supplemented with protease and phosphatase inhibitors (MilliporeSigma). Following incubation on ice for 1 hour, the soluble fraction was separated by centrifugation at 16000 × *g* for 20 minutes at 4^°^C.

### Southern Blot

For Southern blot analysis, 5 micrograms of total cellular DNA was digested with either *Sph*I or *Hind*III, to linearise the HPV16 genome or leave episomes intact, respectively. Digested DNA was separated by electrophoresis using a 0.8% agarose gel, transferred to a nitrocellulose membrane and probed with radiolabeled (32-P) HPV16 genome. This was then visualized by exposure to film for 24 or 72 hours.

### Statistical analysis

Segregation assay results are represented as means ± standard errors (SE). Student’s t test was used to determine significance.

## Acknowledgements

This work was supported by VCU Philips Institute for Oral Health Research and the National Cancer Institute designated Massey Cancer Center grant P30 CA016059 (IMM).

